# Decreased CREB phosphorylation impairs embryonic retinal neurogenesis in the *Oa1-/-* mouse model of Ocular albinism

**DOI:** 10.1101/2024.05.14.594013

**Authors:** Sonia Guha, Andrew M. Nguyen, Alejandra Young, Ethan Mondell, Debora B. Farber

## Abstract

Mutations in the human Ocular albinism type-1 gene *OA1* are associated with abnormal retinal pigment epithelium (RPE) melanogenesis and poor binocular vision resulting from misrouting of ipsilateral retinal ganglion cell (iRGC) axons to the brain. We studied the latter using wild-type (WT) and *Oa1-/-* mouse eyes. At embryonic stages, the WT RPE-specific Oa1 protein signals through cAMP/Epac1-Erk2-CREB. Following CREB phosphorylation, a pCREB gradient extends from the RPE to the differentiating retinal amacrine and RGCs. In contrast to WT, the *Oa1-/-* RPE and ventral ciliary-margin-zone, a niche for iRGCs, express less pCREB while their retinas have a disrupted pCREB gradient, indicating Oa1’s involvement in pCREB maintenance. *Oa1-/-* retinas also show hyperproliferation, enlarged nuclei, reduced differentiation, and fewer newborn amacrine and RGCs than WT retinas. Our results demonstrate that Oa1’s absence leads to reduced binocular vision through a hyperproliferation-associated block in differentiation that impairs neurogenesis. This may affect iRGC axon’s routing to the brain.

## Introduction

Ocular albinism type-1 (OA1) is a developmental eye disease associated with mutations in the *OA1*^1^ gene. It is characterized by two distinct phenotypes: a) abnormal melanogenesis in the retinal pigment epithelium (RPE) where the *OA1* gene is primarily expressed^2^ and b) binocular vision dysfunction^2,3^. The *OA1 g*ene encodes GPR143, a 7-trans-membrane G-protein coupled receptor (GPCR), also known as OA1^4,5^ which directs RPE melanogenesis^6^.

In mice, the Oa1 protein (Gpr143) is first expressed in the RPE at embryonic day (E) 10.5, immediately before melanin is detected at E11^7^. Soon afterwards, ipsilateral retinal ganglion cells (iRGCs) are born between E13.5-E16 in the ventro-temporal (VT) retina; the contralateral (c) RGCs arise throughout the retina from E11 until birth^8^. *Oa1-/-* mice have a reduced number iRGC axons at the optic chiasm than seen in wild-type (WT) mice^2^. This affects binocular vision^3^, the molecular basis of which is unknown. During embryogenesis since the RPE is in physical contact with RGC progenitors near the retinal ciliary margin zones (CMZs), which are areas of progenitor proliferation and migration^9,10^, therefore the RPE may easily send molecular cues to the progenitors that guide early neurogenesis in its immediate vicinity.

The physiological role of the Oa1 protein has been associated in melanocytes with CREB (cAMP response element-binding), a cAMP-responsive transcription factor^11^. In the RPE, the Oa1 GPCR may also interact with the Gα subunit of stimulatory (s) heterotrimeric G-protein(s), to stimulate cAMP synthesis^12^. In general, when a GPCR binds to the Gα_s_ subunit, GDP to GTP exchange occurs on Gα_s_ leading to the dissociation of βγ subunits from the trimeric GTP-Gα_s_. This activates adenylate cyclase and the synthesis of cAMP^12^. cAMP then signals through its effectors: (i) protein kinase A (PKA), which directly phosphorylates (activates) CREB at Ser133^13^ and (ii) EPAC (exchange protein activated by cAMP)^14,15^, which does not act directly on CREB but favors the GDP/GTP exchange by activating the small GTPases Rap1 and Rap2^15,16^. These in turn activate Erk1/2 protein kinases, which phosphorylate CREB^17^. The cAMP-CREB pathway is known to control not only melanogenesis in skin pigmentation^11,18^ but also neurogenesis^19^.

In this study, we explored the possibility of Oa1 being a molecular link between RPE melanogenesis and retinal neurogenesis. Using WT and *Oa1-/-*^20^ mouse eyes, we probed the role of Oa1 in early retinal neurogenesis and axon-routing. Our data indicate that a previously unknown, dynamic Oa1-cAMP/Epac1-Erk2-CREB-dependent activity critically orchestrates neurogenesis at embryonic day (E) 15.5, when RGCs and amacrine cells (ACs) are actively differentiating.

## Results

### Oa1 is exclusively expressed in eyes’s RPE

Figure 1A shows the presence of OA1 protein only in the human adult RPE as previously reported^21^. *In situ* hybridization (RNAscope) corroborates the *Oa1* mRNA exclusive expression in WT E15.5 mouse RPE (Figures 1B and C) and quantitative real-time PCR (*q*RT-PCR) confirms this and shows the absence of *Oa1* mRNA from WT retina. As expected, there is no Oa1 protein expression in either the *Oa1-/-* RPE or retina (Figure 1D).

**Figure 1.**
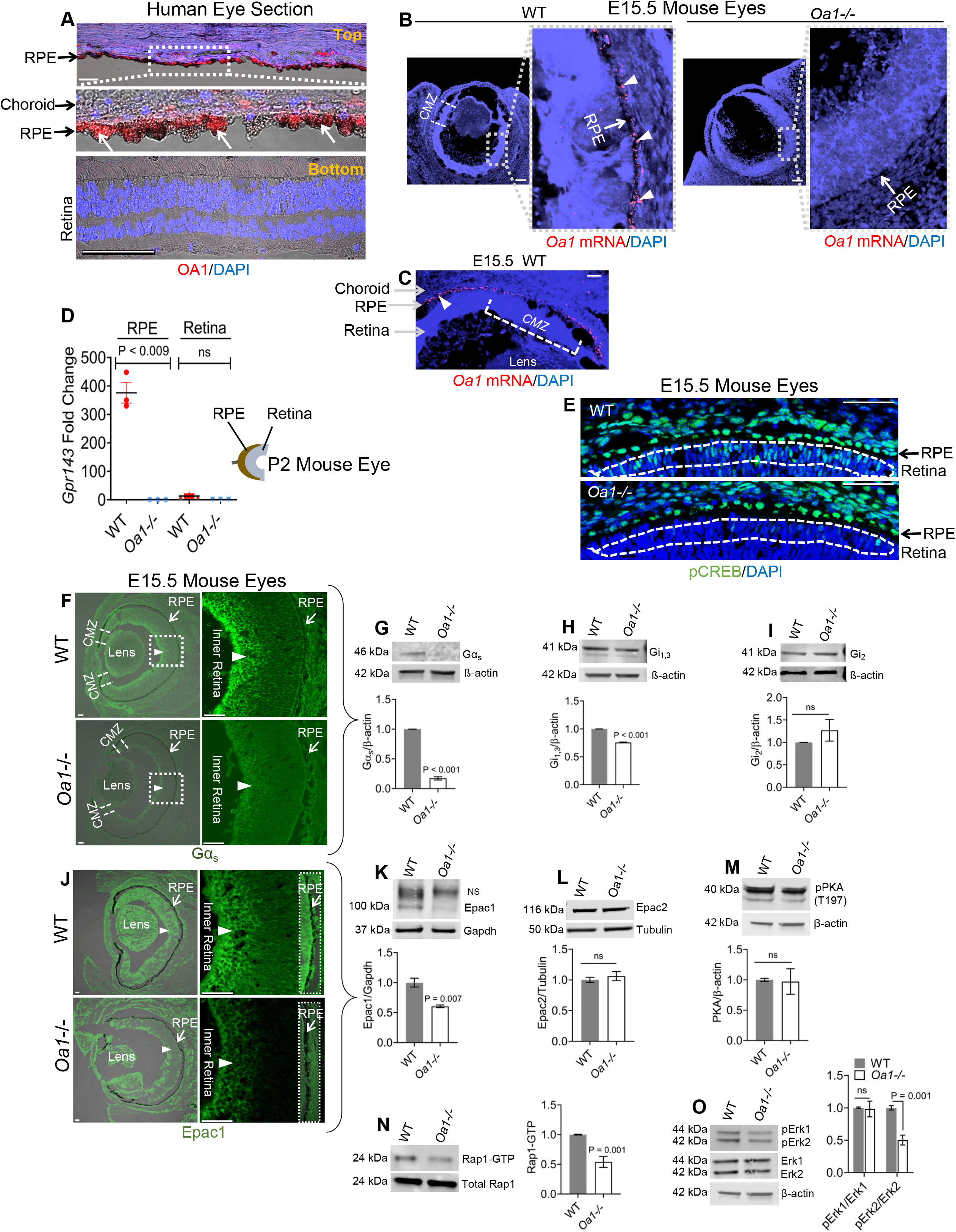
OA1 is only expressed in human and WT mouse RPEs, and its absence is associated with impaired cAMP/Epac1-Erk2 signaling in E15.5 *Oa1-/-* mouse eyes. (A) <u>Top</u>: Immunostaining (red) for OA1 protein in a human donor eye vertical section that has the RPE detached from the retina. The boxed area is magnified below to show prominent staining in the RPE (arrows). <u>Bottom</u>: The retina shows no OA1 staining. DAPI, blue nuclei. Bars, 50 μm. RPE, retinal pigment epithelium. (B-C) Fluorescence *in-situ* hybridization (FISH) with an *Oa1-*specific probe. (B) Magnified boxed regions show intense red punctate staining for *Oa1* mRNA in the WT RPE which is absent in *Oa1-/-* RPE. (C) a different WT eye, showing Oa1 mRNA in the ciliary margin zone (CMZ). Bars, 100 μm. (D) TaqMan *q*RT-PCR of postnatal day 2 (P2) WT and *Oa1-/-* RPEs and retina. RPE and retina are shown sketched. *Oa1* mRNA was normalized to *RPE65* mRNA in RPE and to *ß-actin* mRNA in retina. n=6. (E) IHC with pCREB antibody shows absence of staining in the delineated area of *Oa1-/-* retina compared to WT. Bars, 50 μm. (F) Reduced Gα_s_ staining in *Oa1-/-* inner retina, CMZs and RPE. Boxed regions are magnified on the right. n=3 eyes/group. Bars, 50 μm. (G-I) Western blots show a drastic reduction from WT in the Gα_s_ bands of *Oa1-/-* eyes (G) but not in Gi_1,3_ (H) and Gi2 (I). n=6-9 E15.5 littermates from each of 3 WT and *Oa1-/-* litters. (J) Reduced Epac1 immunostaining in *Oa1-/-* inner retina (arrowheads) and demarcated RPE (arrows). n=3 eyes/group. Bars, 50 μm. (K) Western blots show lower Epac1 band density in *Oa1-/-* eyes. No significant change in Epac2 (L) and pPKA (M). n=6-9 E15.5 littermates, 3 litters/group. (N)Reduced Rap1–GTP levels in *Oa1-/-* eyes after GST–RalGDS–RBD fusion protein pull-down of posterior eyecups. (Rap1-GTP level is a function of Epac1 activity). Bars represent the ratios of Rap1-GTP to total Rap1 in *Oa1-/-* relative to WT eyes. n=5-6 E15.5 litters/group. (O)Western blot shows specific reduction of Erk2 phosphorylation in *Oa1-/-* eyes. n=6-9 E15.5 mice from each litter, 3 litters/group. Significance was determined by Holms-Sidak tests (1A through 1O) except for 1D, analyzed by Welch’s *t-* test.

Transcriptome profiling of RNAs (Microarrays, Affymetrix) isolated from posterior eyecups of E15.5 WT and *Oa1-/-* mice (Figure S1A) shows 51 differentially expressed genes (Figure S1B and Table S1). Out of these, *Gpr143* and four other genes known to be associated with cAMP-signaling are shown in Figure S1C. *q*RT-PCR validated the expression of *Gpr143 (Oa1), Gnas* and *Crebbp* (Figure S1D), and Ingenuity Pathway Analysis (IPA), (Figure S1E) identified CREB as the key transcription factor in this network. *Gnas* and *Crebbp*, which are part of the CREB pathway, are known to be associated with neurogenesis, neuroprotection/survival, and axon pathfinding (Table S2). Based on these data and the known involvement of CREB activation in neuronal differentiation^19^, we followed the presence of phosphorylated CREB (pCREB) in E15.5 eyes; note the sharp difference in pCREB staining next to the RPE between embryonic WT and *Oa1-/-* retinas (Figure 1E, delineated areas).

### cAMP/Epac1-Erk2 signaling is impaired in *Oa1-/-* eyes

The *Gnas* gene encodes Gα_s,_ the alpha*-*subunit of the stimulatory G-protein involved in cAMP production^22^. Immunohistochemistry (IHC) indicates that compared to WT, Gα_s_ is appreciably reduced in the *Oa1-/-* RPEs, CMZs, and inner layers of the peripheral/central retinas (Figure 1F). This is confirmed by western blots that show 6-fold and 1.3-fold decreases respectively, in the levels of Gα_s_ (Figure 1G) and the 41 kDa inhibitory-subunit Gαi_1,3_ (Figure 1H), known to inhibit neural outgrowth^23^.There is no significant change in Gαi_2_ in *Oa1-/-* eyes (Figure 1I).These results suggest that the Oa1-dependent Gα_s_ activation is impaired in *Oa1-/-* eyes, which may impact cAMP production, CREB phosphorylation and its downstream signaling.

We next studied the intracellular cAMP sensors, Epac1 and Epac2, which act as guanine-nucleotide-exchange factors (GEFs) for the small GTPases Rap1 and Rap2, and function in a PKA-independent manner. Epac1 is dominantly expressed in embryonic and neonatal tissues and almost undetectable in the adult^14,24,25^ while Epac2 has an inverted expression pattern^24^. IHC (Figure 1J) shows an appreciable decrease in Epac1 protein in the RPE and retina of E15.5 *Oa1-/-* eyes. Western blots confirm this, with a 1.8-fold reduction from WT in the 100 kDa Epac1 band (Figure 1K) but no discernible change in Epac2 or pPKA (Figures 1L and M).

cAMP induces Epac1’s activity towards its target Rap1^15^. Thus, we investigated if the reduced level of *Oa1-/-* Epac1 would affect Rap1, and through it, the Erk1/2 kinases. A Rap1-GTP pull-down assay with the human Ral GDS-Rap binding domain^26^ demonstrated a 1.84-fold reduction in the activation of *Oa1-/-* Rap1 than WT (Figure 1N), confirming that Oa1 signals through Epac1. This reduced Rap1 activation in turn results in a 2-fold decrease in pErk2 levels but not in pErk1 in *Oa1-/-* eyes (Figure 1O), indicating a role of Epac1 in Erk2 activation^27^.

### Spatio-temporal gradient of pCREB in the developing WT mouse eyes

We examined different areas of the WT RPE and retina (Figure 2A) for pCREB expression at E13.5, E15.5 and postnatal day 2 (P2). Figure 2B shows that at E13.5, pCREB is predominant in the central region (CR) of the RPE/retina, but it is less seen in the dorsal temporal (DT) and ventral temporal (VT) CMZs, which are neurogenic niches; however, their tips show some pCREB expression. At E15.5 and P2, pCREB is detected throughout the RPE and retina. In the CR it forms a temporal gradient that starts in the RPE and progresses through the retinal neuroblasts ending in the differentiating inner retina (Figure 2C). The bar graph on the right of Figure 2C shows this gradient which follows the birth of newborn neurons^28^.

**Figure 2.**
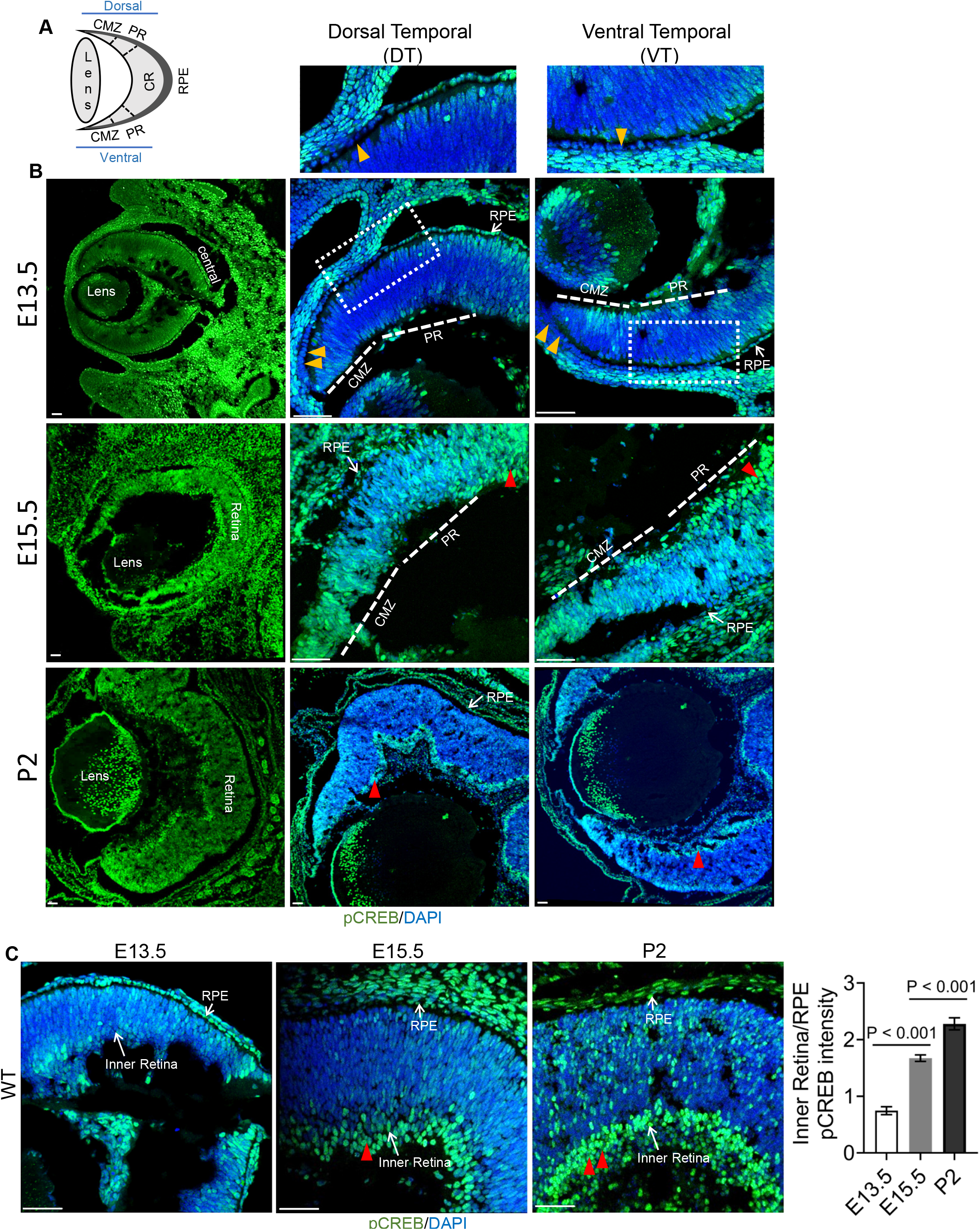
Spatiotemporal expression of pCREB in developing WT mouse eyes. (A)Cartoon^10^ depicting the RPE, CMZs, peripheral retina (PR) and central retina (CR). (B)pCREB expression during development. For each stage, the DT and VT regions are magnified. At E13.5, the images show absence or minimal pCREB in the RPE of the CMZs (yellow arrowheads), PRs and tip of CMZs. pCREB staining increases in E15.5 and P2 retinas (red arrowheads), n=3-4 sections/stage. Bars, 50 μm. (C)pCREB expression in CR of eyes shown in [B]. Quantification showed progressive pCREB increases of 2.2 and 1.4-fold in the inner retina of E15.5 and P2 eyes, respectively, compared to the previous stage (red arrowheads). These eyes were processed together. Significance was determined by Holms-Sidak tests.

### The pCREB gradient is disrupted in *Oa1-/-* eyes

Figure 3A shows pCREB immunoreactivity in E15.5 DT and VT retinal sections of WT and *Oa1-/-* eyes. We only quantified the VT sections since iRGCs arise from the vCMZ^10^. Compared to WT, pCREB intensity is lower by 40% and 45% in the vCMZ and peripheral retina (PR) of *Oa1-/-* eyes, respectively, (Figure 3A, bar graph). In Figure 3B, the concurrent pCREB staining intensity decrease in the *Oa1-/-* CR RPE (38%) and stark absence in the developing *Oa1-/-* neuroblasts (*) suggests that the RPE significantly contributes to disrupt the pCREB gradient. This is supported by western blots of E15.5 posterior eyecup lysates (Figure 3C) that similarly detects a 2 and 1.6-fold reduction in the respective band densities of *Oa1-/-* pCREB and pATF1, a CREB-related gene^29^. However, at P2, pCREB is only decresed 1.5-fold from WT and there is no change in pATF1 in *Oa1-/-* eyes (Figure 3D). We also found that overexpressing *in vitro* the OA1 protein in ARPE-19 cells (a cell line derived from human RPE) enhances their pCREB staining (Figures S2A and B).

**Figure 3.**
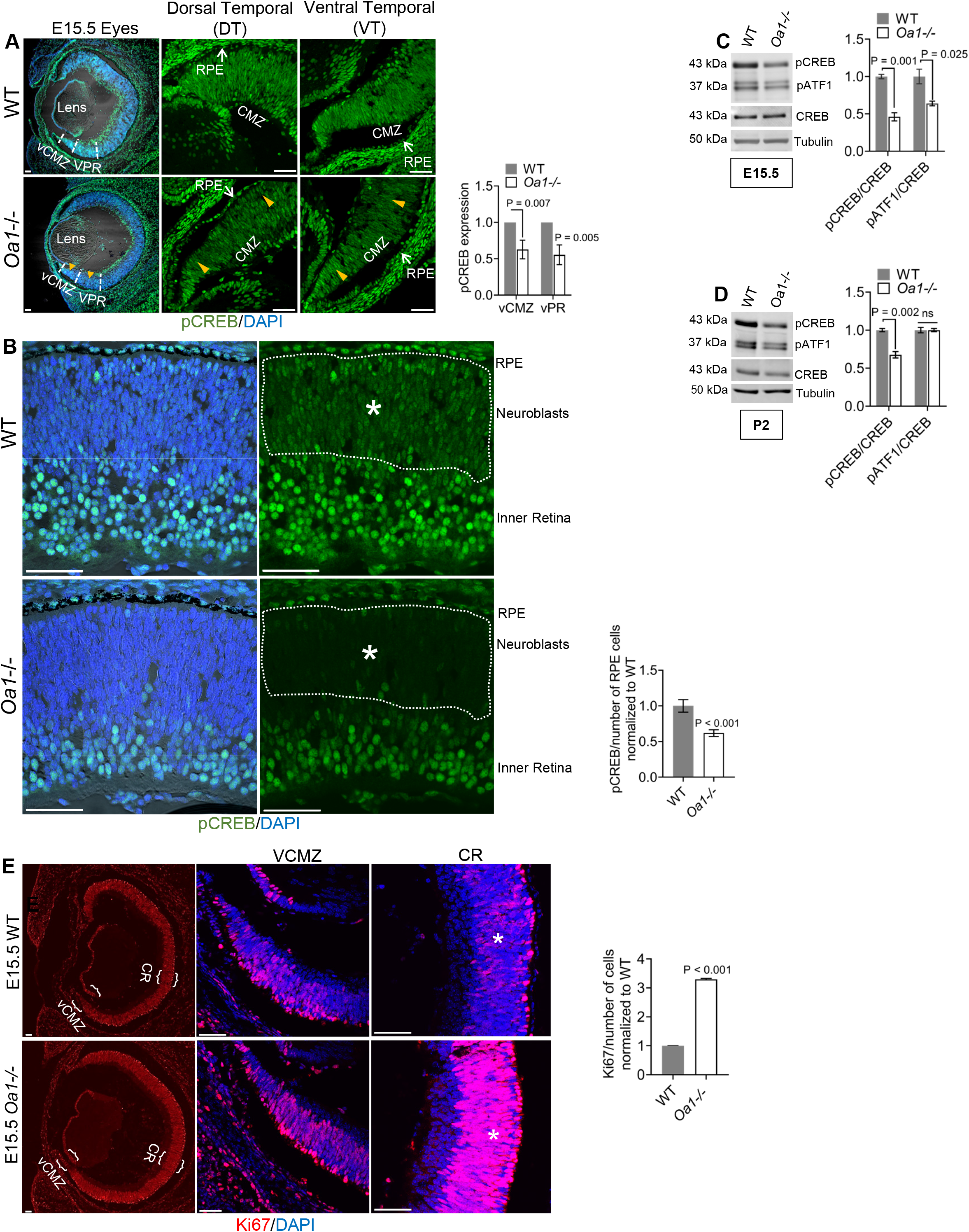
Reduced pCREB expression and retinal hyperproliferation in *Oa1-/-* eyes. (A)Representative images show reduced pCREB expression in *Oa1-/-* eyes (arrowheads). DT and VT regions are zoomed on the right. These eyes were processed together. Bar graphs show reduced pCREB expression in *Oa1-/-* vCMZ and vPR (demarcated with white-dashed lines on whole eyes), n=3-6 E15.5 eyes/group. vCMZ, ventral ciliary margin zone; ventral peripheral retina (vPR). Bars, 50 μm. (B)Images from the CR of the eyes in [A] show reduced pCREB staining in *Oa1-/-* RPE, neuroblasts (asterisks) and inner retina. Bar graph shows reduced pCREB in *Oa1-/-* RPE. (C-D) Western blots and bar graphs show reduced pCREB and pATF1 levels in *Oa1-/-* eyes. n=3-12 littermates, 3 litters/group. (E) Demarcated regions from the CR and vCMZ in WT and *Oa1-/-* eyes are magnified to the right. IHC of vCMZ and CR and quantification of the CR (asterisks) show hyperproliferation (enhanced Ki67 staining) in *Oa1-/-* eyes. Bars, 50 μm Significance was determined by Holms-Sidak tests.

Collectively, these data establish the presence of an Oa1-induced Epac1-Erk2-CREB signaling cascade in embryonic mouse RPE/retina. Intravitreal injection of the EPAC inhibitor ESI-09^30^ or the ERK inhibitor FR180204^31^ in 2-months-old WT mouse eyes disrupts this cascade, providing further support for it. Conversely, an Epac1 agonist, 8CPT^32^ completely rescues the ESI-09-blocked Epac1 protein levels and simultaneously increases the levels of pErk2, pCREB and pATF1 proteins. These data demonstrate that Epac1 signals downstream to Erk2 and CREB in mouse eyes and that Epac1-Erk2-CREB are arranged as serial components in a single pathway (Figure S3).

### pCREB retinal immunoreactivity expands from E13.5 to E15.5 in WT eyes

At E13.5, mostly non-proliferating early progenitors (Ki67-/Nestin+) express pCREB. These cells are clustered around the optic stalk and in the CMZs (Figures S4A and B, E13.5). With development, at E15.5, pCREB is detected in the CMZs that now harbor both proliferating (Ki67+) and non-proliferating progenitors (Ki67-) (Figures S4A and B, E15.5), in the newly differentiated cells of the forming RGC layer (Ki67-) and in the highly proliferative cells near the RPE-retina contacts (Ki67+, demarcated Figure S4B, E15.5). Neurogenesis also follows a similar pattern and increases after E13.5^8,33^.

A select population of pCREB+ cells in the vCMZ of E15.5 WT eyes also express the cell-cycle exit marker cyclin D2 (Figure S4C) reflecting pCREB’s role in cell-cycle exit^34^ and iRGC neurogenesis^10^. These cells are non-proliferating (Figure S4D), pointing to pCREB’s expression by early iRGC progenitors in the CMZ. Both non-proliferating and proliferating cells in the CMZs and near the RPE-retina region are in contact with Oa1-positive RPEs, as shown above (Figure 1B and C). This, together with the fact that E15.5 RPE-retina contact areas and vCMZs have reduced pCREB intensity in *Oa1-/-* eyes (Figures 1E and 3A), suggests that Oa1-CREB signaling at RPE-RGC contacts is disrupted during embryogenesis.

### Enhanced proliferation and reduced differentiation in *Oa1-/-* retinas

Surprisingly, a comparative assessment of expression of the cell-cycle marker Ki67 between E15.5 WT and *Oa1-/-* eyes, revealed a 3.3-fold increase in the latter (Figure 3E), indicating hyperproliferation^35^. Since cell proliferation and differentiation show a remarkable inverse relationship, we next assessed the expression of Tuj1, a neuron-specific class III *β*-tubulin. Tuj1 is a marker for differentiating immature neurons^36^. Consistent with pCREB’s role in cell differentiation^36^, *Oa1-/-* retinal cells also show decreases from WT of 3% in intensity and 46% in number of Tuj1+ cells/unit area (Figure 4A), suggesting a hyperproliferation-associated differentiation-block in E15.5 *Oa1-/-* eyes.

**Figure 4.**
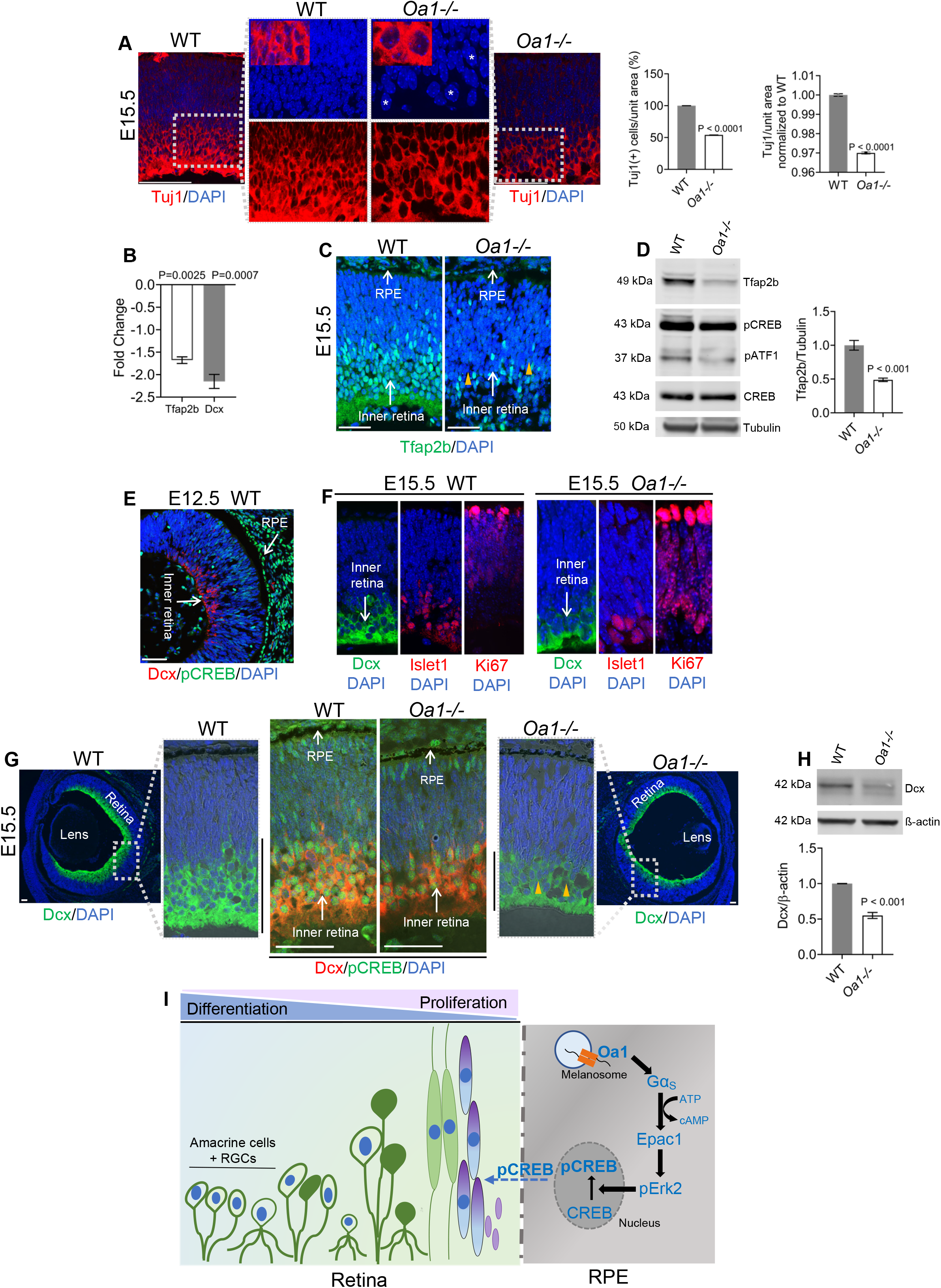
Impaired AC and RGC growth in embryonic *Oa1-/-* eyes. (A) Images show *Oa1-/-* retina has larger nuclei (asterisks), fewer Tuj1+ cells and reduced Tuj1 intensity per unit area, respectively. The boxed areas show zoomed retinal cells. Bars, 50 μm. (B) qRT-PCR showing reduced *Tfap2b and Dcx* mRNAs in E15.5 *Oa1-/-* eyes compared to WT. (C) Reduced Tfap2b (AC marker) expression in E15.5 *Oa1-/-* mouse eyes and retinas (yellow arrowheads). Bars, 50 μm. (D) Western blot and bar graphs show reduced Tfap2b level in *Oa1-/-* eyes. pCREB and pATF1 levels are also reduced in *Oa1-/-* eyes. n=6-12 E15.5 littermates, 3 litters/group. (E) Representative image shows Dcx+ RGCs (red) in the central WT retina. (F) Dcx+ RGCs from E15.5 retinas are non-proliferating (Ki67-) and express Islet1, a marker of differentiated RGCs. Ki67 staining is in a different section than that shown in [Figure 3E]. (G) E15.5 eyes show an expansion in Dcx (green) staining. The zoomed images (orientation changed) show reduction of Dcx in *Oa1-/-* mouse retinas (yellow arrowheads). The length of the Dcx labeled neurites (black line bar) is shorter in *Oa1-/-* retinas. Co-labelling for Dcx (red) and pCREB (green) shows both markers decreased in *Oa1-/-* eyes. DAPI, nuclei. (H) Western blot and bar graphs show reduced Dcx level in E15.5 *Oa1-/-* eyes. n=6-12 littermates, 3 litters/group. Significance was determined by Holms-Sidak tests, except in 4A and 4B, analyzed by Welch’s *t-*tests. (I) Schematic representation of Oa1-induced CREB activation in an embryonic mouse eye. RPE-specific Oa1 signals mainly through Epac1-pErk2 to phosphorylate CREB in the RPE. This is followed by CREB activation in the adjacent retina, leading to proliferation and differentiation.

### Decrease of newborn neurons in *Oa1-/-* eyes

We next checked for retinal neural markers which appear when the cells exit the cell-cycle and differentiate. Table S1 shows that two key retinal neural markers, (Tfap2b and Dcx), have lower expression in E15.5 *Oa1-/-* than in WT eyes. Tfap2b is a transcription factor belonging to the Activating Enhancer Binding Protein 2 family. It is transcriptionally co-activated by Crebbp^37^ and facilitates AC differentiation during retinogenesis^38^. *q*RT-PCR and IHC show a 1.7-fold reduction from WT in the *Oa1-/-Tfap2b* transcript (Figure 4B) and a drastic decrease of Tfap2b-stained nuclei in the inner retina (Figure 4C), respectively. Western blots confirmed these results, with a 2-fold decrease in *Oa1-/-* Tfap2b band density in E15.5 eyes (Figure 4D).

Doublecortin (Dcx) is a microtubule associated protein required for initial steps of neuronal migration^39^. We detected Dcx in pCREB positive cells of E12.5 WT eyes localized mainly in the CR (Figure 4E), consistent with a previous study showing that the first RGCs that extend to the brain arise from the CR at E12^40^. In line with the role of CREB phosphorylation in neuronal differentiation and doublecortin expression^41,42^, we show that E15.5 mouse eyes express pCREB in the differentiated non-proliferating RGCs that are positive for Islet1 (Figure 4F), as reported earlier for E14 mouse retina^43^. At this stage, both pCREB and Dcx expression expand from the center of the retina towards the PR (Figure 4G). Notably, the length of the Dcx-stained neurites is shorter in *Oa1-/-* eyes, pointing to a reduced migration of retinal progenitors (Figure 4G). *q*RT*-*PCR and western blot data likewise confirm 2 and 1.8-fold reduction in *Dcx* transcript and protein levels, respectively, in *Oa1-/-* E15.5 eyes (Figures 4B and H) indicating compromised neurogenesis.

## Discussion

Previously, we had shown that mutations in the *Oa1* gene causally lead to Ocular albinism, a disease of abnormal melanogenesis in the RPE^2^. In this study, we demonstrate that the Oa1 protein, an RPE-specific 7-transmembrane-GPCR controls early neurogenesis in mouse embryonic eyes by regulating the activation of the cAMP-dependent transcription factor CREB, an intermediate in GPCR-activated dopamine signaling^44^.

Oa1-coupled Gα_S_ produces cAMP, which in turn activates Epac1. In developing mouse eyecups, absence of Oa1 decreases Epac1 expression resulting in reduced levels (Figures 1J, K and N) of Epac-induced Rap1-GTP^45^. In neuronal cells, Oa1 activates Erk^23^ and Epac1-Erk1/2 signaling induces axonal growth via CREB activation^27^. CREB functions are potentiated by ATF1; both act in concert during early embryogenesis^29^ to recruit co-activators like Crebbp^46^ and transactivate an array of CREB-dependent genes^47^. We report impaired phosphorylation of Erk2, CREB and ATF1 along with reduced Crebbp levels in *Oa1-/-* eyes (Figures 1O, 3C and S1D). The robust CREB phosphorylation dynamics seen in the RPE, the adjoining neuroblast layer and the differentiating inner retina of WT eyes is markedly disrupted in *Oa1-/-* eyes, remaining almost undetected in their neuroblasts (Figures 1E and 3B). Furthermore, the decrease in Oa1-associated CREB activation in *Oa1-/-* RPE (Figure 3B) is responsible for the substantial disruption of the pCREB gradient seen in *Oa1-/-* eyes. Out of the many pathways that can activate CREB^48^, our data highlights the key role of Oa1 in CREB-dependent signaling during early mouse retinogenesis and underlines the impact of an RPE-specific gene activity in the birth of retinal neurons.

CREB regulates cell-cycle exit^34^. In agreement with this, E15.5 *Oa1-/-* retinal cells are arrested in their cell-cycle (Figure 3E). Based on our data, we propose that CREB activation by Oa1 regulates the birth of retinal cRGCs from E11-to birth, and it directs the appearance of iRGCs in the vCMZ, by E15. During this critical window, CREB probably influences the expression of cyclin D2 in the iRGC progenitors of the vCMZ (Figures S4C and D), enabling them to exit the cell-cycle and to differentiate into neurons. Diminished CREB activation in the vCMZ of E15.5 *Oa1-/-* eyes (Figure 3A) possibly alters cell-cycle exit and leads to a reduction in iRGCs^10^.

CREB-deficient embryos have developmental eye defects^19^. We show that Oa1 significantly impacts CREB activity in WT E15.5 eyes, when the RGCs are actively growing, in comparison to the developing eyes (Figure 3D) just after birth, when their RGCs have already crossed the chiasm^8^. Therefore, absence of Oa1 at E15.5 affects CREB-dependent RGC-genesis, routing, and the neural wiring in the optic-chiasm responsible for the binocular circuit and depth perception.

Collectively, our results show that the absence of Oa1 is associated with impaired CREB phosphorylation, increased proliferation, decreased differentiation, enlarged cell size and thereby fewer retinal cells. Such CREB activity-related differentiation-block in *Oa1-/-* RGCs and ACs (Figures 3 and 4) during the peak of retinal neurogenesis at E15.5, may compromise visual signal processing, as seen in albinos^49^.

Based on our findings, we speculate that after its expression in mouse RPE at E10.5^7^, Oa1 activates the cAMP-signaling cascade that we are describing in embryonic RPE (Figure 4I). This possibly influences microtubule-based melanosome motility and explains why melanosomes are distributed towards the periphery in *Oa1-/-* RPEs as early as E15.5^50,51^. Alternatively, Oa1-cAMP-CREB may activate the RPE-specific *Mitf* isoforms A, D, H and M^52^ to impact melanogenesis, similar to what is reported for mouse melanocytes^11^. In addition, since the RPE makes physical contact with the CMZs and the outer retina, CREB activation could also initiate RGC neurogenesis. Infact, activation of CREB in the RGC progenitors of the CMZs and the neural retina is supported by the detection of a pCREB gradient in the developing retina (Figure 2C). Thus, CREB could be a key regulator of both RPE melanogenesis and RGC routing.

Clinically, patients carrying mutations in the *Oa1* gene have giant melanosomes in their RPEs and poor binocular vision. The data presented in this paper suggests that the two apparently ‘unrelated’ phenotypes (melanogenesis in the RPE and neurogenesis in the retina) may share a common molecular link, the Oa1-cAMP signaling cascade that activates CREB. By integrating embryonic melanogenesis and neurogenesis this cascade may explain why albinism is associated with photo aversion, decreased visual acuity, and impaired depth perception.

## Limitations of the study

Although to the best of our knowledge this is the first report to show the critical role of an RPE-expressed protein, Oa1, in early retinal neurogenesis, future work is needed to identify the factors that drive it. For example: Does the RPE-expressed pCREB communicate with the retina directly, or indirectly through gap-junctions, notch signaling or RPE secreted CREB-dependent neurotrophins known to influence retinal CREB phosphorylation and neurogenesis^53-55^? Neither did we determine whether the Oa1-Epac pathway is completely independent or if it cooperates with PKA^56^ and related mediators like AKAPs^57^ during early retinogenesis.

## Acknowledgments

We thank Dr. Roxana Radu (UCLA) for providing the human eye sections and Dr. Suraj Bhat (UCLA) for his comments on the manuscript. Yisha He helped with the experiments. This work was supported by a Vision of Children Foundation grant (S.G.) and an NIH Center Core grant P30EY000331 (D.B.F.).

## Author Contributions

S.G., and D.B.F. designed this study. S.G. procured grant funding. S.G., A.M.N., and E.M. performed the experiments and analyzed the results. A.Y., injected the animals. S.G., wrote the paper. S.G., and D.B.F. edited the manuscript.

## Declaration of Interests

The authors declare no competing interests.

## METHODS

### Data and code availability

Microarray dataset is available under GEO accession number GSE174287. Any additional information required to reanalyze the data reported in this work paper is available from the Lead contact upon request.

## EXPERIMENTAL MODEL AND SUBJECT DETAILS

### Animals

#### Mice

C57BL/6NCrl (WT) and congenic *Oa1* knockout mice (*Oa1-/-*) were obtained from The Charles River Labs (USA and Italy, respectively) and bred at UCLA. The Murine Genetic Analysis Laboratory, UC Davis, confirmed that the *Oa1-/-* mice were congenic using the methodology previously described ^2^. For timed mating, mice were housed in a timed-pregnancy breeding colony at UCLA. In both colonies, females were checked for vaginal plugs at approximately noon each day. E0.5 corresponds to the day when the vaginal plug was detected, with the assumption that conception took place at approximately midnight. Our studies were carried out according to protocols approved by the UCLA Animal Research Committee and the ARVO Statement for Use of Animals in Ophthalmic and Vision Research. 2-6 months-old male and female mice were used for mating.

#### Cell line culture

The immortalized human RPE cell line ARPE-19 cells (ATCC) were grown to confluence in 75 cm^2^ culture flasks in a 1:1 mixture of Dulbecco’s modified Eagle medium and Ham’s F12 medium with 100 U/mL penicillin, 100 μg/mL streptomycin, plus 10% fetal bovine serum (Thermofisher).

## METHOD DETAILS

### Embryo collection

Six pregnant females (three *Oa1-/-* and three WT), were euthanized at E13.5 or E15.5. Eyes from six embryos of each type were removed and treated as one biological sample. After enucleation, the anterior segments were dissected out and the posterior eyecups were stored at -80°C for Western blots, and overnight at 4°C in RNAlater (Thermofisher) for RNA isolation. Theiler Stage guidelines were used for the phenotypical characterization of embryonic eyes.

### Isolation of RPE and retina from WT and *Oa1-/-* postnatal day 2 (P2) pups and of posterior eyecups from adult WT mice

P2 pups and adult mice were euthanized by hypothermia or isoflurane inhalation, respectively, followed by cervical dislocation. After enucleation, the anterior segments of pup eyes were quickly removed and the retinas dissected, washed in Tris-buffered saline (TBS) and stored at -80°C. The RPE was then gently separated from the eyecup using a loop and stored at -80°C. For adult mice, following enucleation and removal of anterior segments, the posterior eyecups were dissected, washed in TBS and stored at -80°C.

### RNA isolation

Total RNA from each biological sample was isolated using the RNAqueous-4RT-PCR kit (Thermofisher) and then treated with 2 units of amplification grade DNase I (Thermofisher). The RNA concentration was quantified, and its quality assessed using a Nanodrop ND-1000 spectrophotometer (Thermofisher) and also by electrophoresis on agarose gels and staining with ethidium bromide. 18S and 28S RNA bands were visualized under ultraviolet light. An absorbance ratio of 260/280 measured in a bioanalyzer (Agilent) was used to assess RNA purity.

### Microarray hybridization and data analysis

Total RNA from E15.5 posterior eyecups (retina, RPE, choroid and sclera) of each of 3 *Oa1-/-* and 3 WT mice was isolated as described above and reverse transcribed into cDNA following the manufacturer’s instructions. Each of the 6 cDNAs was hybridized in triplicate to the 430.20 Affymetrix Mouse Microarrays at the UCLA Clinical Microarray Facility. Microarray results were gathered using the Affymetrix Gene Chip Operating Software and sample data was normalized to WT to determine log ratios between diseased and WT samples. Differences in gene expression levels between *Oa1-/-* and WT samples with p-values < 0.05 and a fold change ≥ 1.5 were considered statistically significant. Raw data were analyzed for obtaining PCA maps using Partek Genomics Suite 6.4 software. Global functional analysis, network analysis and canonical pathway analysis were performed using Ingenuity Pathway Analysis (IPA, Ingenuity Systems).

### Quantitative real time PCR (qRT-PCR) validation of microarray results

The sequences of differentially expressed genes were obtained using BLAST genomic databases (NCBI) and two different sets of mouse forward and reverse primers were designed from them. After optimizing the primer conditions using Primer 3 Output (NCBI), the best set of primers for each gene showing good dissociation curves and no primer dimerization was selected as follows. *Gpr143*(*Oa1*): forward 5’-TGGTGATTCAGTGGGAAACA; reverse 5’-AGGACCCAGTTGCAGTATGG; *Gnas:* forward 5’-CTCATCGACAAGCAACTGGA; reverse 5’-CCTGCACTTTAGTGGCCTTC; *Crebbp*: forward 5’-GTCTTTGCCTTTTCGTCAGC; reverse 5’-CCACATACTGCCAGGGTTCT; *Dcx*: forward 5’-TCGTAGTTTTGATGCGTTGC; reverse 5’-ACAGACCAGTTGGGGTTGAC; *Tfap2b*: forward 5’-CCCGGGCCGTTTATCTCTAC; reverse 5’-CCCGCGGGTAAATTCAAACC; *B-actin:* forward 5’-CCTAAGGCCAACCGTGAAAAGATG; reverse 5’-ACCGCTCGTTGCCAATAGTGATG. The master mix for qRT-PCR contained 2.5 μl of Brilliant SYBR green 2X (Agilent), 1μl of 150 nM forward and reverse primers, 0.4 μl of diluted reference dye (1:500) and 10.1 μl of water to complete a final volume of 24 μl per reaction. 1μl of cDNA per sample was added in triplicate to the plate and the fully loaded plate was gently mixed, centrifuged and run using a Mastercycler RealPlex (Eppendorf). The cycling parameters were set as follows: 10 min at 95ºC, then 40 cycles of denaturation at 95°C for 30 sec, followed by annealing for 1 min at 60°C and elongation at 72°C for 30 sec. Three different qRT-PCR experiments were carried out using the RNA sample aliquots from the microarrays. All samples were run in triplicates. In each experiment, a sample without reverse transcriptase and a sample without template were included to demonstrate specificity and lack of DNA contamination.

Following the qRT-PCR experiment, gene differences for each sample on the 96 well-plate were quantified using cycle threshold (Ct) values (number of cycles required for the fluorescent signal to exceed background data level). Ct values for *Oa1-/-* and WT samples were used to compare gene expression data (ΔCT) for each gene of interest (GOI). Expression of genes in *Oa1-/-* samples relative to that of a housekeeping gene (HKG), ΔCT_*Oa1-/-* GOI_ = CT_*Oa1-/-* GOI_ – CT_HKG,_ was compared to the expression of the same gene in WT samples, ΔCT_WT_ = CT _WT-GOI_ - CT_HKG_. The relative gene expression level (fold change in the *Oa1-/-* sample over that in the WT sample) was calculated using (2^-ΔCt^) for each gene in *Oa1-/-* and in WT samples and their ratio was determined.

### qRT-PCR analyses of Oa1 expression in RPE and retina of P2 mice

qRT-PCR was performed using Taqman 2X Universal PCR Master Mix, No AmpErase UNG, and Taqman probes (Thermofisher) specific for Oa1 (Mm00440553_m1), or for the endogenous controls RPE65 (Mm00504133_m1) and ß-Actin (Mm02619580_g1). The qRT-PCR master mix contained 18 ul of 2X Universal PCR Master Mix, 1 μl of Taqman probe and 1 μl of cDNA (100 ng) per sample. All samples were run in triplicates. Thermal cycling parameters were set up for 1) 10 min at 95°C, 2) 15 sec at 95°C, and 3) 60 sec at 60°C, repeating steps 2 and 3 for 40 cycles.

### RNAscope® based Fluorescent In-situ hybridization (FISH)

E15.5 mouse heads were fixed in 4% paraformaldehyde in 0.1M phosphate buffer (PB), pH 7.4 for 24 hours at 4°C, then washed in 0.1M phosphate-buffer saline (PBS), followed by sequential incubation in 10% and 30% sucrose for cryoprotection. The heads were then embedded in cryo-OCT (Tissue-Tek), cut into 10 μm sections, mounted onto Super-Frost Plus slides and stored at -80°C for future use. *In-situ* hybridization was performed on the fixed frozen tissue sections at the UCLA Translational Pathology Core laboratory using the automated RNAscope LS Multiplex Fluorescent Assay [Advanced Cell Diagnostics, (ACD), catalog #32280] of the Leica Biosystems BOND RX platform, according to the manufacturer’s protocol. Prior to the automated procedure, the slides were washed in 1X PBS to remove the OCT, baked for 30 min at 60ºC and post-fixed in 4% PFA in 1X PBS for 15 min at 4ºC. The tissues were then dehydrated using sequentially 50, 70 and 100% ethanol. Slides were air dried 5 min at room temperature (RT). During the automated RNAscope assay the tissues were hybridized with the following appropriate ACD probes: mouse *Gpr143* (catalog *#*535108-C2), positive probes (mouse *Polr2A*-C1/*Ppib*-C2/*Ubc*-C3, catalog #320888) and negative probes (bacterial *dapB*-C1/*dapB*-C2/*dapB*-C3, catalog #320878). Fluorescent signals were detected by the Opal fluorophores (Opal 520, 570 or 690; Akoya Biosciences). The tissues were mounted with ProLong Gold Antifade Mountant (ThermoFisher) and imaged with the Vectra Polaris Imaging System (Akoya Biosciences).

### Western Blotting

Posterior eyecups from *Oa1-/-* and WT E15.5 and P2 pups, and from adult WT mice that had been stored at -80°C were homogenized manually with a cordless grinder (Kimble-Chase) in Tissue Protein Extraction Reagent (T-PER,Thermofisher) with protease and phosphatase inhibitor cocktail (Thermofisher), followed by centrifugation at 12,000g for 10 min at 4°C. Protein concentrations were determined using the BCA kit (Thermofisher). Samples were mixed with 4X NuPAGE LDS sample buffer (Thermofisher), heated at 70°C for 3 min and proteins were separated under non-reducing conditions by either 7% or 4-12% Bis-Tris SDS-polyacrylamide gel electrophoresis (PAGE) using 1X MOPS/SDS running buffer, pH 7.0 (Bioland Scientific). The separated proteins were transferred to PVDF membranes (Bio-Rad) overnight. After treating the membranes with Odyssey blocking buffer (Li-COR Biosciences) they were incubated with the appropriate primary antibodies listed below, overnight, at 4°C. The primary antibodies were diluted with 1% Tween20 in 1X Tris-buffered saline (TBS): Odyssey®TBS Blocking Buffer (1:1). The membranes were then washed three times (15 min each) with TBS containing 2% Tween20 and incubated with anti-rabbit or anti-mouse fluorescently tagged (IRDye) secondary antibodies for 1 hr at RT. Bands were captured and quantified using an Odyssey Imaging System (Li-COR, Image Studio Lite Version 5.2.5). Equal protein loading was confirmed with antibodies against ß-actin, tubulin or Gapdh, as appropriate.

List of primary antibodies used: Rabbit anti-PKA (1:5000; Abcam, Cat#ab75991), Rabbit anti-Gα_s_ (1:1000; Abcam, Cat#ab83735), Rabbit anti-Gαi_1,3_ (1:1000; Abcam, Cat#ab154024), Mouse anti-Gαi_2_ (1:1000; SCBT, Cat#sc-13534), Mouse anti-Epac1(1:500; SCBT, Cat#sc-28366), Rabbit anti-Epac1(1:1000, Abcam Cat#ab109415), Rabbit anti-Epac2 (1:1000; Proteintech, Cat#19103-1-AP), Rabbit anti-phospho CREB (1:1000, Cell Signaling, Cat#9198), Mouse anti-CREB (1:1000, Cell Signaling, Cat#9104), Rabbit anti-phospho Erk1/2 (1:1000, Cell Signaling, Cat#9101), Rabbit anti-Erk1/2 (1:1000, Cell Signaling, Cat#9102), Rabbit anti-Dcx (1:2000, Abcam, Cat#ab6142), Rabbit anti-AP-2beta (1:1000, Cell Signaling, Cat#2509), Rabbit anti-Gapdh (1:2000, Abcam, Cat#ab9485), Mouse anti-ß-actin (1:2000, Abcam, Cat#ab8226), Mouse anti-alpha-tubulin (1:5000, Abcam, Cat#ab4074).

### Immunohistochemistry (IHC)

Mouse heads of E12.5, E13.5, E15.5 and P2 pups or posterior eyecups from adult eyes were immersed in 4% paraformaldehyde in 0.1M phosphate-buffer (PB), pH 7.4, overnight, at 4°C. Next, they were washed with three changes of PB, transferred to 10% sucrose in PB and incubated for 2 hr at 4°C, followed by 30% sucrose in PB for 4 hr at 4°C and finally by 1:1 30% sucrose: O.C.T. tissue embedding compound (Tissue Tek) overnight, at 4°C. The fixed heads or posterior eyecups were then frozen in liquid nitrogen and stored at -80°C. When ready to use, they were placed for 30 min in a -20°C freezer and then 10-um thick vertical sections were cut on a freezing microtome (Leica). Before immunostaining, the sections were blocked in either 10% non-specific goat serum (NGS) or donkey serum (NDS) in PBS for 1 hr at RT to prevent non-specific binding. Sections were then incubated overnight at 4°C with the appropriate primary antibodies listed below. After washing three times in PBS containing 0.5% Triton X-100, the sections were incubated for 1 hr with the corresponding Alexa Fluor-conjugated-secondary antibodies. Sections were then washed several times with PBS containing 0.5% Triton X-100, stained with 4’, 6-diamidino-2-phenylindole (DAPI) and mounted in Fluoromount-G medium (Southern Biotech). Confocal images were acquired using a FluoView FV1000 confocal laser-scanning microscope (Olympus). Images were processed using Adobe Photoshop software. Projected 3D images were for 21 z-positions; n = 3 sections per group. The confocal images were processed with the IMARIS software (Bitplane) or Adobe Photoshop software.

List of primary antibodies used: Rabbit anti-GPR143 (1:500, Sigma, Cat#HPA003648), Rabbit anti-Gα_s_ (1:100; Abcam, Cat#ab83735), Rabbit anti-Epac1(1:100, Abcam, Cat#ab21236), Rabbit anti-phospho CREB (1:100, Cell Signaling, Cat#9198), Rabbit anti-phospho Erk1/2 (1:100, Cell Signaling, Cat#9101), Rabbit anti-Dcx (1:200, Abcam, Cat#ab6142), Rabbit anti-AP-2beta (1:1200, Cell Signaling, Cat#2509), Rabbit anti-Ki67 (1:1000, Abcam, Cat#ab15580), Rabbit anti-Islet1 (1: 500, Abcam, Cat#ab20670), Mouse anti-Nestin (1:100, Abcam, Cat#ab6142), Mouse anti-Tuj1 (1:500, Abcam, Cat#ab78078), Mouse anti-CyclinD2 (1:50, SCBT, Cat#sc-56305).

### Immunocytochemistry (ICC)

Cells were grown on coverslips and were washed with PBS for 20 min at RT, permeabilized in ice cold methanol at −20°C for 15 min and subsequently blocked in 0.3% TritonX-100 and 3% NDS at RT for 1hr, followed by overnight incubation in primary antibodies, Mouse anti-GPR143 (1:50, SCBT, Cat#sc-398602) and Rabbit anti-phospho CREB (1:100, Cell Signaling, Cat#9198) at 4°C. Next day, cells were washed three times in PBS containing 0.5% Triton X-100 and incubated in Alexa Fluor-conjugated secondary antibodies for 1 hr at RT. Coverslips were then washed, mounted onto the glass slides using ProLong Gold Antifade Mountant with DAPI (ThermoFisher) and imaged using a FluoView FV1000 confocal laser-scanning microscope.

### Intravitreal injection of inhibitors/activator

EPAC1 inhibitor ESI-09 (Tocris) and ERK1/2 inhibitor FR-80204 (Millipore Calbiochem) were dissolved in dimethylsulfoxide (DMSO) to get 3.3 mM stock solutions. These stocks were subsequently diluted to the final 1μM concentration using 1X PBS. A 20 mM stock solution of EPAC activator, 8CPT-2Me-cAMP (8CPT, Tocris), was prepared in phosphate-buffered saline (PBS) and subsequently diluted to 20.5 μM final concentration. Briefly, the pupil of the anesthetized eye was dilated with tropicamide drops. 1 μl of the appropriate inhibitor (1 μM) or activator (20.5 μM) was injected into the vitreous through a post limbus spot using a Hamilton microinjector under a stereoscopic microscope (Carl Zeiss). At the appropriate time points, animals were sacrificed and those with visible lens damage or vitreal hemorrhage were excluded from the analysis.

### ARPE19 OA1-GFP Stable Transfection

#### Expression construct

A pCMV6-Entry plasmid containing human *OA1* cDNA open reading frame with a C-terminal GFP tag was purchased from OriGene technologies (Catalog # RG209560). This plasmid has the complete CMV promoter and ampicillin and geneticin resistance for selection.

#### Stable transfection and selection of stably transfected ARPE19 cells

The day before transfection, each well of three 6-well plates was seeded with 8x10^5^ ARPE19 cells (50-80% confluent). The ARPE19 cells were then transfected with the *OA1*-GFP construct using the PrimeFect1 DNA transfection reagent (Lonza), according to the manufacturer’s protocol. Prior to selection of transfected cells using 200 μg/ml geneticin (Gibco), both transfected and non-transfected control cells were incubated at 37ºC for 72 hours prior. Cells were allowed to recover from exposure to the transfection reagents or from resuspension and re-plating, after passage of confluent cultures for one day in complete medium without antibiotics. Selected cells were plated on T75 culture flasks and incubated in a humidified incubator at 37°C, 5% CO_2_. Fresh complete or selection medium containing geneticin was added daily and morphology of cells was monitored by microscopy.

### Quantification and statistical analysis

Sample sizes, statistical tests, and P values are indicated in the figure legends. All statistical analyses were performed using Prism version 8.02 (GraphPad). The values obtained are indicated as mean ± SEM. The brightness and contrast of confocal images were adjusted using an Olympus FV10-ASW (Version 04.2a). Measurements were done blinded to mouse genotypes.

For quantification of pCREB signal intensity in the vCMZ and vPR, analysis were performed on at least 3 different samples per age and genotype. At least 10 sections were taken from each sample. Pixel intensity of pCREB+ cells was measured with ImageJ analysis software and the intensities of the *Oa1-/-* sections were normalized to those in the WT sections. For spatiotemporal quantification, the ratio of pCREB intensity in the inner retina to that in the RPE of a WT mouse eye at different developmental stages was determined. For quantification in the RPE, the ratio of pCREB intensity in the RPE to the number of DAPI-stained RPE cells was measured and *Oa1-/-* values were normalized to WT set as 1. For quantification of Ki67+ cells in the E15.5 retinas, a defined area in the CR was analyzed and at least 10 sections from 3 animals of each genotype were used. The intensity of Ki67 measured by ImageJ was divided by the number of cells and the *Oa1-/-* values are normalized to WT, set as 1. For quantitation of cell differentiation, the number of Tuj1-stained cells in a defined region of the CR of E15.5 *Oa1-/-* eyes was analyzed by ImageJ and reported as percent of WT or with respect to that in WT set as 1. The Tuj1 intensity in this region was also measured with respect to WT, set as 1. 10 sections from 3 animals of each genotype were analyzed.

## Notes

### Competing Interest Statement

The authors have declared no competing interest.

