## Supplemental Files for "Decreased CREB phosphorylation impairs embryonic retinal neurogenesis in the *Oa1-/-* mouse model of Ocular albinism"

### Figure S1. Transcriptomic analysis of E15.5 eyes from WT and *Oa1-/-* mice

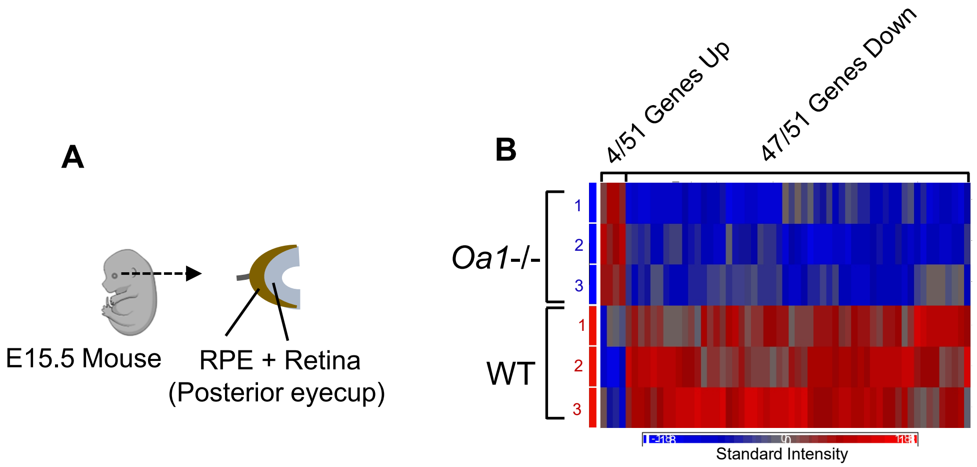


**C**

| Gene Symbol | Gene Name | p-value | Fold Change |
|---|---|---|---|
| ***Gpr143*** | **G protein-coupled receptor 143** | **0.0001** | **-5.791** |
| ***Gnas*** | **GNAS (guanine nucleotide binding protein, alpha stimulating) complex locus** | **0.0092** | **-1.992** |
| *Akap9* | A kinase anchor protein (yotiao) 9 | 0.0058 | -1.885 |
| ***Crebbp*** | **CREB binding protein** | **0.0427** | **-1.665** |
| *Rif1* | Rap1 interacting factor homolog | 0.0446 | -1.517 |

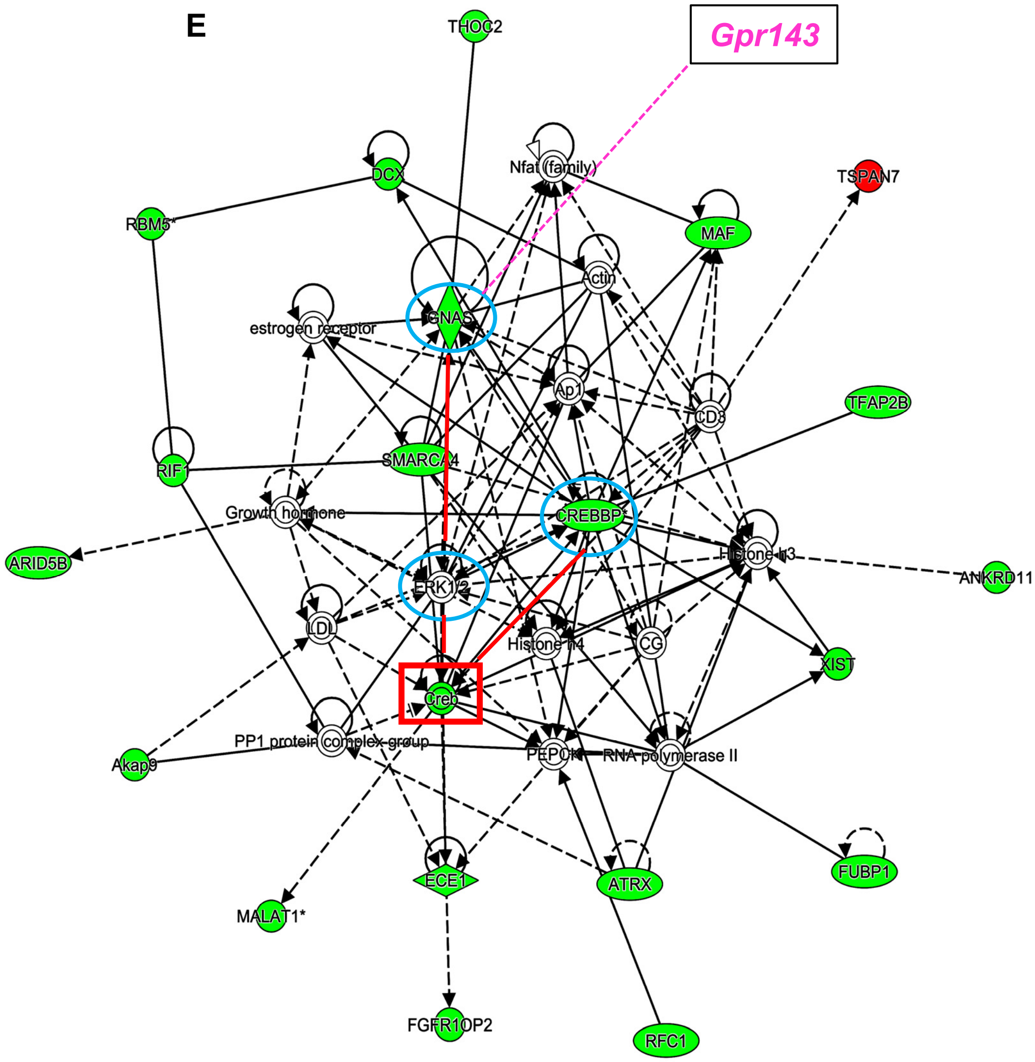


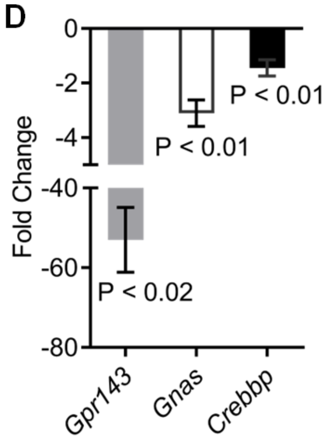

**Figure S2. OA1 increases CREB phosphorylation in human RPE cells**

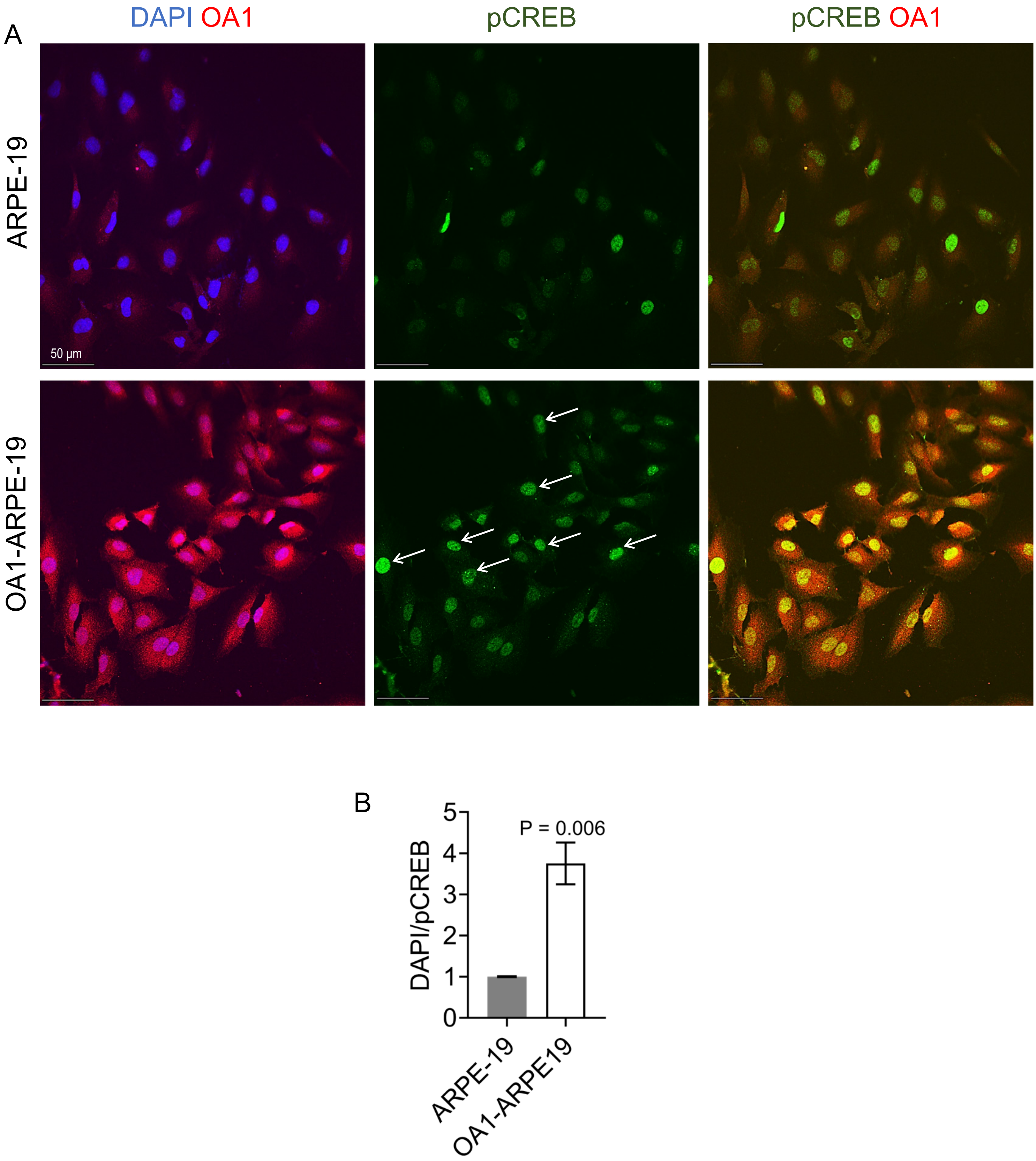

#### Figure S3. cAMP-Epac1-Erk2-CREB signaling in WT mouse eyes

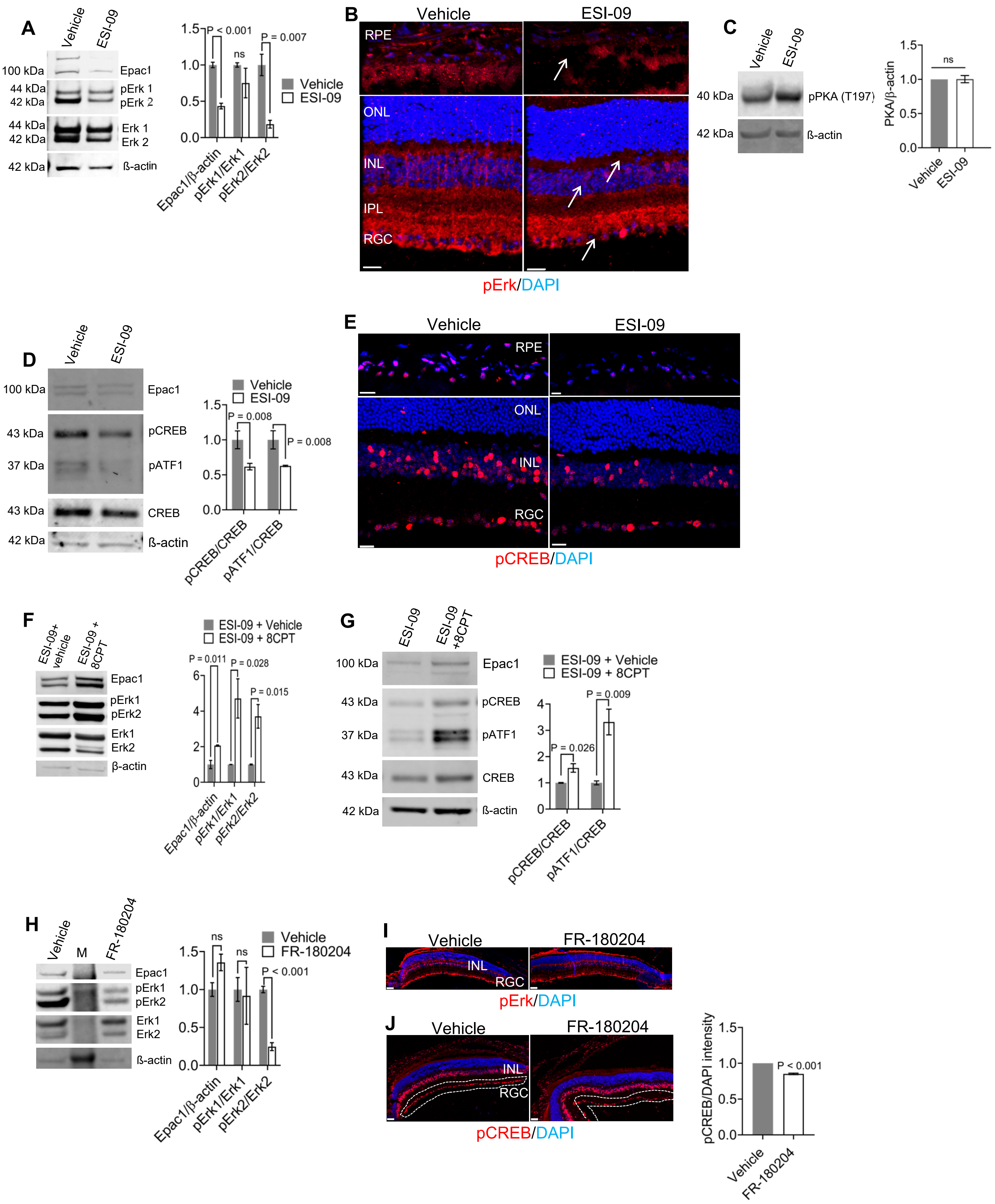

#### Figure S4. pCREB is expressed by non-proliferating, proliferating progenitors and differentiated retinal cells

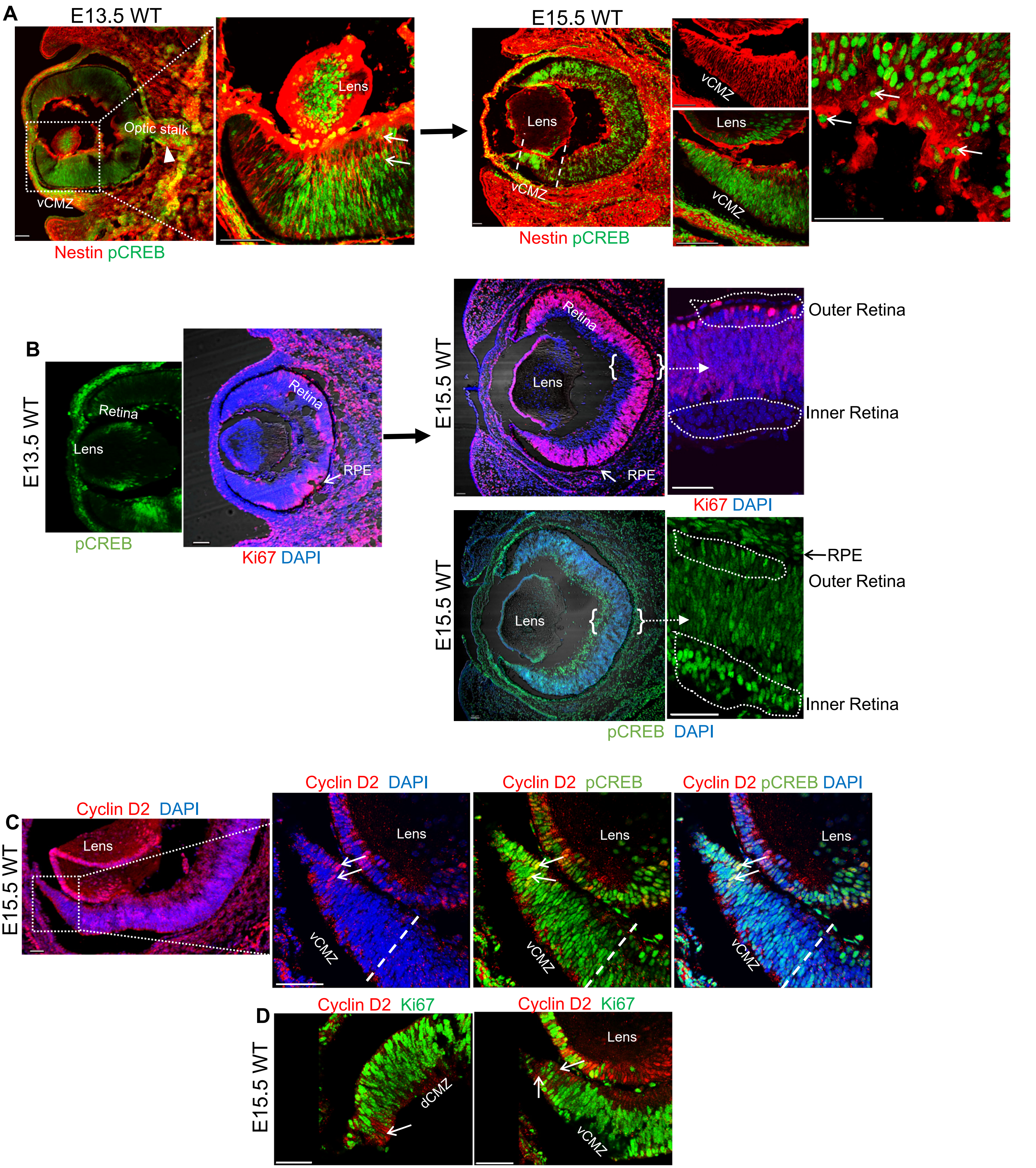

**SUPPLEMENTAL INFORMATION TITLES AND LEGENDS**

**Figure S1. Transcriptomic analysis of E15.5 mice eyes. Related to Figure 1.**

(A) Total RNAs extracted from E15.5 WT and *Oa1-/-* posterior mouse eyecups were subjected to microarray analysis. (B) Of the 51 differentially expressed genes seen in the Heat map representation of the results, 4 genes had increased (red) and 47 decreased (blue) expression in *Oa1-/-* eyes compared to the WT, with a fold-change (FC) ≥1.5 and a p-value <0.05); rows = samples (1,2,3, biological repeats); columns = genes.

(C) Differentially expressed genes in *Oa1-/-* and WT eyes affecting the cAMP pathway[1-5] (D) qRT-PCR validation of the differential expression of *Gpr143, Gnas and Crebbp* from [C]. A bar graph shows the transcripts fold-change in *Oa1-/-* samples compared to WT normalized to 1. n=3 RNA samples tested in each group. Significance was determined by Welch's t-test. (E) Ingenuity pathway analysis (IPA) identifies CREB (red square) as a key transcription factor in the *Gpr143* gene network. Green = downregulated and red = upregulated genes. Genes not colored were not identified in our microarrays but are relevant to this network. The genes we have studied are encircled in blue and connected by red lines. The pink dashed line shows the possible link between *Gpr143* and the gene network.

**Figure S2. Increased CREB phosphorylation in OA1 transfected human RPE cells. Related to Figures 2 and 3.**

(A) Representative images of human RPE cells, ARPE-19, immunostained with antibodies that recognize pCREB (green) and OA1 (red). The upper panel shows non-transfected ARPE-19 cells while the lower panel has cells transfected with OA1-expressing plasmid. DAPI (blue) shows cells nuclei. (B) The number of pCREB stained nuclei increases by 3.7-fold in OA1 transfected cells. The bar graph shows quantification of the ratio of fluorescence intensity of pCREB-stained nuclei to the same nuclei stained with DAPI and normalized to non-transfected cells set as1. Significance was determined by Holms-Sidak test.

**Figure S3. cAMP-Epac1-Erk2-CREB signaling in WT mouse eyes. Related to Figures 1 and 3.**

(A-C) Epac-mediated Erk activation. (A) Western blot of posterior eyecup lysates from 2-months old WT mice 72 hrs after intravitreal injection of the ESI-09, an Epac inhibitor (1µM), shows decreased intensity of Epac1 and pErk2 bands compared to vehicle-injected. pErk1 did not change significantly. These results indicate an impaired Epac1-mediated Erk2 phosphorylation. The bar graph depicts the ratios of band density for Epac1 normalized to ß-actin and pErk1/2 normalized to Erk1/2, respectively, in vehicle (set as 1) and ESI-09-injected eyes. n=3 mice per group. ESI-09 binds to the functional cAMP-binding pocket of Epac1. The Epac1 antibody used here targets sequences in the C-terminus catalytic region of Epac1. (B) A section from an ESI-09-injected eye shows reduced pErk staining (red) indicated by arrows than sections from a vehicle injected eye. n=3 sections per group; DAPI (blue) shows nuclear staining. ONL, outer nuclear layer; INL, inner nuclear layer; IPL, inner plexiform layer; RGC, retinal ganglion cell. Bar, 20 µm. (C) There is no difference in phosphorylated PKA (pPKA) levels of vehicle and ESI-09 injected eyes.

(D-E) Epac1 induced CREB activation. (D) Western blots of posterior eyecup lysates of mice intravitreally injected with vehicle or ESI-09 indicate that ESI-09 effectively blocks Epac1, pCREB, and pATF1. pCREB and pATF1 bands quantified and normalized to CREB are shown relative to those in vehicle-injected eyes set as 1 (n=3). (E) Immunostaining for pCREB in eyes of 2-months old WT mice injected with ESI-09 shows reduced signal in RPE and retina when compared to vehicle-injected eyes. Bar, 50 µm.

(F-G) 8CPT rescues Erk and CREB activation. (F) 72 hrs after ESI-09 treatment, 2-months-old WT eyes were either re-injected with vehicle or 8CPT (20.5 μM). Western blots of these eye lysates show restoration of Epac1, pErk1/2, and Erk1/2 levels 5 days after 8CPT re-injection. Bar graphs represent the ratios of band density for Epac1 to ß-actin or pErk1/2 to Erk1/2, respectively, in ESI-09+8CPT-treated samples normalized to ESI-09-injected eyes set as 1. n=3 mice per group. Note that 8CPT rescued both Epac1 and pErk1/2 levels. (G) Other WT eyes injected with 20.5 μM, 8CPT 72 hrs after ESI-09 injection also show restored levels of pCREB and pATF1 when compared to vehicle-injected eyes. pCREB/CREB and pATF1/CREB signal ratios were quantified and normalized to ESI-09-injected eyes, n=3 mice per group.
(H-J) FR-180204, an Erk inhibitor, blocks pErk and pCREB signals indicating Erk-induced CREB phosphorylation. (H) Western blots of posterior eyecup lysates of 2-months-old WT mice intravitreally injected with vehicle or FR-180204 show a significant difference in the density of pErk2 but not Epac1 or pErk1 bands 72 hrs after injecting FR-180204 (n=3). Epac1 and pErk1/2 signals were quantified and normalized to ß-actin and Erk1/2, respectively, and are plotted relative to vehicle-injected eyes. M, molecular weight markers. (I) Immunostaining for pErk1/2 (red) shows FR-180204 blocked pErk expression in the INL and RGCs of WT mice (n=3 sections per group). Bar, 50 μm. (J) Immunostaining for pCREB (red) in sections from mice eyes injected with vehicle or FR-180204 shows reduced pCREB staining of the RCG layer in FR-180204 injected eyes, also indicating that inhibition of Erk2 signaling affects CREB phosphorylation. Bar, 50 μm. Intensity of pCREB/DAPI stained nuclei for the RGC layers (demarcated by dotted lines) is plotted normalized to vehicle injected eyes (n=3 sections per group).
Significance was determined by Holms-Sidak test for all the experiments carried out in Figure S3.

**Figure. S4. pCREB is expressed by non-proliferating, proliferating progenitors and differentiated retinal cells. Related to Figures 3 and 4.**
(A) Representative images of WT eye sections show that pCREB is expressed by retinal progenitors (nestin+) during early embryogenesis. Left panels (of E13.5 and E15.5 eyes) show expression of both pCREB (green) and nestin (red) around the optic stalk (arrowhead in E13.5) and in the vCMZs at E13.5 and E15.5. The boxed areas having the vCMZs are magnified at the right of the whole eyes. pCREB and nestin co-expression is indicated by white arrows. n=3 WT eye sections. Bar, 50 μm.
(B) pCREB is expressed by proliferating Ki67+ cells in E13.5 and in E15.5 (near the RPE-retina contact, demarcated in the zoomed image) eye sections. It is also expressed by differentiated retinal cells (Ki67-, demarcated with dotted lines) in the zoomed E15.5 section on the right. n=3 WT eye sections. Bar, 50 μm. DAPI (blue) shows nuclear staining.
(C)The boxed area is magnified on the three panels to the right to show co-localization of cyclin D2 with pCREB (yellow, in the middle and the right panel indicated by white arrows) in the vCMZ. n=3 E15.5 WT eye sections. Bar, 50 μm.
(D) Cyclin D2 is expressed by non-proliferating (Ki67-) progenitors in the CMZs, pointed by white arrows. n=3 E15.5 WT eye sections. Bar, 50 μm. dCMZ, dorsal CMZ; vCMZ, ventral cMZ.

**Table S1. Differential gene expression from E15.5 eyes in WT vs *Oa1-/-* mice.**

**Table S2. Functions of selected down-regulated genes identified in E15.5 *Oa1-/-* mice eyes.**

**Table. *S*1.** Differential gene expression from E15.5 eyes in WT vs *Oa1-/-* mice

| Gene Symbol | Gene Name | RefSeq Transcript ID | p-value | Fold-Change |
|---|---|---|---|---|
| Upregulated genes at E15.5 | | | | |
| Pisd-ps3 | phosphatidylserine decarboxylase, pseudogene 3 | NR_003518 | **0.0426883** | **3.34934** |
| Snrpn// Snurf | small nuclear ribonucleoprotein N /// SNRPN upstream reading frame | NM_001082961/// NM_001082962/// NM_013670 /// NM_033174 | **0.0353932** | **1.70437** |
| Tspan7 | tetraspanin 7 | NM_019634 | **0.0469085** | **1.58658** |
| Defa15 | defensin, alpha, 15 | --- | **0.0109935** | **1.51158** |
| Downregulated genes at E15.5 | | | | |
| Sfrs18 | serine/arginine-rich splicing factor 18 | NM_025669 | **0.022383** | **-1.5063** |
| Thoc2 | THO complex 2 | NM_001033422 | **0.01905** | **-1.5099** |
| Rif1 | Rap1 interacting factor 1 homolog (yeast) | NM_175238 | **0.044674** | **-1.5173** |
| Arid5b | AT rich interactive domain 5B (MRF1-like) | NM_023598 | **0.001227** | **-1.5196** |
| Nipbl | Nipped-B homolog (Drosophila) | NM_027707/// NM_201232 | **0.036951** | **-1.5241** |
| Dclre1a | DNA cross-link repair 1A, PSO2 homolog (S. cerevisiae) | NM_018831 | **0.002** | **-1.5252** |
| Smarca4 | SWI/SNF related, matrix associated, actin dependent regulator of chromatin, subf | NM_001174078/// NM_001174079/// NM_011417 | **0.023612** | **-1.5317** |
| Birc6 | baculoviral IAP repeat-containing protein 6 | NM_007566 | **0.046478** | **-1.5344** |
| Scaper | S phase cyclin A-associated protein in the ER | NM_001081341/// NM_175536 | **0.019907** | **-1.5358** |
| Baz1b | bromodomain adjacent to zinc finger domain, 1B | NM_011714 | **0.003342** | **-1.5362** |
| Maf | avian musculoaponeurotic fibrosarcoma (v-maf) AS42 oncogene homolog | NM_001025577 | **0.043078** | **-1.5367** |
| 2900092E17Rik/ // Prrt2 | RIKEN cDNA 2900092E17 gene/// proline-rich transmembrane protein 2 | NM_001102563/// NM_030240 | **0.033343** | **-1.5546** |
| Tmem26 | transmembrane protein 26 | NM_177794 | **0.039703** | **-1.5457** |
| Ccdc58 | coiled-coil domain containing 58 | NM_001159421/// NM_001159422/// NM_198645 | **0.047402** | **-1.5496** |

| Gene Symbol | Gene Name | RefSeq Transcript ID | p-value | Fold-Change |
|---|---|---|---|---|
| Tfap2b | transcription factor AP-2 beta | NM_001025305/// NM_009334 | **0.000539** | **-1.5546** |
| Atrx | Alpha thalassemia/mental retardation syndrome X-linked homolog (human) | NM_009530 | **0.014591** | **-1.5611** |
| Ece1 | endothelin converting enzyme 1 | NM_199307 | **0.04268** | **-1.5721** |
| Zfp398 | zinc finger protein 398 | NM_027477 /// NM_173034 | **0.025329** | **-1.5723** |
| Prrx1 | paired related homeobox 1 | NM_001025570 /// NM_011127 /// NM_175686 | **0.026162** | **-1.5806** |
| Mll5 | myeloid/lymphoid or mixed-lineage leukemia 5 | NM_026984 | **0.00122** | **-1.5826** |
| C77673 | expressed sequence C77673 | --- | **0.036725** | **-1.6074** |
| Zer1 | zer-1 homolog (C. elegans) | NM_178694 | **0.013042** | **-1.6307** |
| Gbf1 | golgi-specific brefeldin A-resistance factor 1 | NM_178930 | **0.042264** | **-1.6479** |
| Rbmx | RNA binding motif protein, X chromosome | NM_001166623/// NM_011252 /// NR_029425 | **0.010826** | **-1.663** |
| Crebbp | CREB binding protein | NM_001025432 | **0.042657** | **-1.6646** |
| Eif3a | eukaryotic translation initiation factor 3, subunit A | NM_010123 | **0.01161** | **-1.6791** |
| Mia3 | melanoma inhibitory activity 3 | NM_177389 | **0.008376** | **-1.686** |
| Etnk1 | ethanolamine kinase 1 | NM_029250 | **0.034037** | **-1.6983** |
| Gtf3c1 | general transcription factor III C 1 | NM_207239 | **0.024564** | **-1.6992** |
| Ankrd11 | ankyrin repeat domain 11 | NM_001081379 /// NR_037865 | **0.003518** | **-1.7058** |
| Naa15 | N(alpha)-acetyltransferase 15, NatA auxiliary subunit | NM_053089 | **0.041788** | **-1.713** |

| Gene Symbol | Gene Name | RefSeq Transcript ID | p-value | Fold-Change |
|---|---|---|---|---|
| 5830407P18Rik | RIKEN cDNA 5830407P18 gene | --- | **0.013023** | **-1.7667** |
| Rfc1 | replication factor C (activator 1) subunit 1 | NM_011258 | **0.01179** | **-1.7817** |
| Prrc2c | proline-rich coiled-coil 2C | NM_001081290 | **0.020661** | **-1.793** |
| Fgfr1op2 | FGFR1 oncogene partner 2 | NM_026218 | **0.020785** | **-1.8306** |
| Erdr1 | erythroid differentiation regulator 1 | NM_133362/// XR_141421 /// XR_142424 | **0.007563** | **-1.8722** |
| Akap9 | A kinase (PRKA) anchor protein (yotiao) 9 | NM_194462 | **0.005875** | **-1.885** |
| Fubp1 | Far upstream element (FUSE) binding protein 1 | NM_057172 | **0.0198349** | **-1.96926** |
| Gnas | GNAS (guanine nucleotide binding protein, alpha stimulating) complex locus | NM_001077507///NM_001077510 ///NM_010309///NM_010310/// NM_019690 /// NM_0 | **0.00917** | **-1.9901** |
| Malat1 | Metastasis associated lung adenocarcinoma transcript 1 (non-coding RNA) | NR_002847 | **0.044307** | **-2.0588** |
| Zc3h15 | zinc finger CCCH-type containing 15 | NM_026934 | **0.00867977** | **-2.03526** |
| Rhox4b | Reproductive homeobox 4B | NM_021300 | **0.00570379** | **-2.06588** |
| Rbm5 | RNA binding motif protein 5 | NM_148930 | **0.00883866** | **-2.19024** |
| Dcx | Doublecortin | NM_001110222///NM_001110223/// NM_001110224 /// NM_010025 | **0.003049** | **-2.5926** |
| Xist | inactive X specific transcripts | NR_001463 /// NR_001570 | **0.028026** | **-3.1584** |
| Gpr143 | G protein-coupled receptor 143 | NM_010951 | **0.000107** | **-5.7905** |
| 2900056M20Rik | RIKEN cDNA 2900056M20 gene | NR_040269 /// NR_040270 | **0.000622** | **-5.815** |

**Table S2.** Functions of selected down-regulated genes identified in E15.5 *Oa1-/-* mice eyes

| Gene Symbol | Gene Name | Function |
|---|---|---|
| *GPR143* | G protein-coupled receptor 143 | Embryonic Development, Developmental Disorder, G-Protein Coupled Receptor Signaling, Visual System Development and Function, Ophthalmic disease, Hereditary disorder, Tissue Morphology, Cell Morphology, Cellular Assembly and Organization, Cell Signaling[1,2]. |
| *GNAS* | GNAS (guanine nucleotide binding protein, alpha stimulating) | Embryonic Development, Developmental Disorder, Tissue Morphology, Molecular Transport, Protein Synthesis, Small Molecule Biology, Cell Signaling, Cell Cycle[3,4]. |
| *CREBBP* | CREB binding protein | Embryonic Development, Developmental Disorder, Nervous System Development and Function, Neurological Disease, Tissue Morphology, Cellular Development, Cell Morphology, Cell Signaling, Cellular Function and Maintenance[5,6]. |
